# Bacterial Polyphosphates Induce CXCL4 and Synergize with Complement Anaphylatoxin C5a in Lung Injury

**DOI:** 10.1101/2022.07.05.498838

**Authors:** Julian Roewe, Sarah Walachowski, Arjun Sharma, Kayleigh A. Berthiaume, Christoph Reinhardt, Markus Bosmann

## Abstract

Polyphosphates are linear polymers of inorganic phosphates that exist in all living cells and serve pleiotropic functions. Bacteria produce long-chain polyphosphates, which can interfere with host defense to infection. In contrast, short-chain polyphosphates are released from platelet dense granules and bind to the chemokine, CXCL4.

Here, we report that long-chain polyphosphates induced the release of CXCL4 from mouse bone marrow-derived macrophages and peritoneal macrophages in a dose-/time-dependent fashion resulting from an induction of CXCL4 mRNA. This polyphosphate effect was lost after pre-incubation with recombinant exopolyphosphatase (PPX) Fc fusion protein, demonstrating the potency of long chains over monophosphates and ambient cations. In detail, polyphosphate chains longer than 70 inorganic phosphate residues were typically required to mediate robust CXCL4 release. Polyphosphates acted independently of the purinergic P2Y1 receptor and the MyD88 and TRIF adaptors of Toll-like receptors. On the other hand, polyphosphates augmented LPS/MyD88-induced CXCL4 release, which was explained by intracellular signaling convergence on PI3K/Akt. Polyphosphates alone induced phosphorylation of Akt at threonine-308. Pharmacologic blockade of PI3K (wortmannin, LY294002) antagonized polyphosphate-induced CXCL4 release from macrophages. Intra-tracheal polyphosphate administration to C57BL/6J mice caused histologic signs of lung injury, disruption of the endothelial-epithelial barrier, influx of Ly6G^+^ polymorphonuclear neutrophils, depletion of CD11c^+^SiglecF^+^ alveolar macrophages, and release of CXCL4. Long-chain polyphosphates synergized with the complement anaphylatoxin, C5a, which was partly explained by upregulation of the receptor, C5aR1, on myeloid cells. C5aR1^-/-^ mice were protected from polyphosphate-induced lung injury. In conclusion, we demonstrate that polyphosphates govern immunomodulation in macrophages and are capable of inducing lung injury.

## Introduction

Inorganic polyphosphates (PolyP) are polymers of phosphate residues joined by high-energy phosphoanhydride bonds, which may be as short as three or as long as 1,700 residues *in vivo* (1, 2). Polyphosphates are ubiquitous and evolutionarily conserved, and previous studies have characterized both regulatory and chaperone functions beyond their original roles as phosphate and energy reservoirs (1, 3, 4). For a homopolymeric molecule, polyphosphates affect an astounding number of biological functions, including metabolic regulation through stimulation of mTOR (5), virulence and resistance to oxidative stress (6), regulation of complement activation (7), and blood clotting with polymer length-dependent effects in humans (2). Platelets typically produce polyphosphates with a size of less than 100 monomers long, whereas bacteria produce polyphosphates of several hundreds or more units (2).

Polyphosphates interfere with the differentiation and function of macrophages during bacterial infection (8-10). Polyphosphates prevent phagosome acidification and thus enhance survival of bacteria after phagocytosis by macrophages (8, 10). The MHCII-dependent antigen presentation machinery is suppressed by polyphosphates (8). LPS-primed macrophage polarization is skewed towards an M2-resembling phenotype characterized by suppression of NOS2 as a marker for M1 macrophages (8, 11). In addition, long-chain polyphosphates reduce LPS-induced STAT1 phosphorylation that is required for type I interferon production (8). Accordingly, the production of the type I interferon induced chemokine, CXCL10, is decreased by long-chain polyphosphates (8). The phagocytosis-induced release of CCL2 is blocked in macrophages (8). All these effects are specific to long-chain polyphosphates, which overwhelmingly antagonize beneficial host responses. On the other hand, short-chain polyphosphates (the length produced by platelets *in vivo*) enhance CXCL4 binding to *E. coli* and trigger neoepitope anti-CXCL4/polyanion antibody-mediated phagocytosis by polymorphonuclear neutrophils (PMNs) (12).

CXCL4, also known as platelet factor 4 (PF4), is a chemokine which binds polyanionic molecules like polyphosphates and others such as heparin, DNA, RNA, and lipid A (12). CXCL4 ligates with the CXCR3 receptor in mice and the CXCR3B splice variant in humans (13). CXCL4 plays regulatory roles in hemostasis/thrombosis, inflammation, angiogenesis and wound healing (14, 15).

Bacterial infections can lead to the life-threatening complications such as sepsis, acute lung injury (ALI), and acute respiratory distress syndrome (ARDS). ALI/ARDS are severe respiratory conditions characterized by pulmonary edema, inflammation of the alveolar epithelium, and destruction of healthy lung tissue (16). ALI/ARDS and sepsis often co-exist in the same critically ill patients and have limited FDA-approved treatment options. We have recently reported that long-chain polyphosphates mediate death in *E. coli* sepsis and that targeting polyphosphates using recombinant exopolyphosphatase increases survival in mice (8). However, a pathogenetic role of long-chain polyphosphates in ALI/ARDS has not yet been investigated. In contrast, numerous studies over recent decades have highlighted the critical role of the anaphylatoxin, C5a, in the pathogenesis of ALI/ARDS (17-19). C5a is a potent peptide generated during complement activation, which binds the C5aR1 receptor to directly promote chemotaxis of PMNs and induce chemokines in myeloid cells (16). Intratracheal C5a administration to healthy mice is sufficient to unleash a robust inflammatory response similar to human ARDS (16).

Here, we identified the property of bacterial-type, long-chain polyphosphates to induce CXCL4 release from macrophages and during lung injury. The acute deleterious effects of polyphosphates in the lung were chain-length dependent and amplified by complement component C5a rather than by CXCL4.

## Methods

### Mice

All animal experiments were approved by the State Investigation Office of Rhineland-Palatinate and the Institutional Animal Care and Use Committee (IACUC) of Boston University. All mice were housed in a 45-60% humidity, 22±2°C ambient temperature, controlled light/dark (12h/12h) cycle, with free access to food and water. C57BL/6J mice (8-12 weeks old), C5aR1^-/-^ mice, MYD88^-/-^ mice, TRIF^-/-^ mice, and P2Y1^-/-^ mice (all on C57BL/6J background) were maintained in a specific pathogen-free environment. Sex-age matched cohorts were used for experiments.

### Macrophages

Bone marrow-derived macrophages (BMDMs) were generated by culturing mouse bone marrow cells with L-929 cell-conditioned medium for 7 days (18). Peritoneal-elicited macrophages (PEMs) were isolated 4 days after i.p. injection of 1.5 ml 2.4% (w/v) thioglycollate (Becton Dickinson, Franklin Lakes, NJ). The cultivation of macrophages was performed in RPMI 1640 (Thermo Fisher Scientific) supplemented with 100 U/ml penicillin-streptomycin (Thermo Fisher Scientific) and 0.1% (w/v) bovine serum albumin (Carl Roth, Karlsruhe, Germany) at 37°C, 5% CO_2_, and 95% humidity.

### Lung Injury

Mice were anesthetized with ketamine/xylazine before surgical exposure of the trachea on an upright surgical stand (17). The following substances were slowly injected intra-tracheally (i.t.) in 40 μl phosphate buffered saline (PBS): Synthetic polyphosphates of the doses and chain-lengths as indicated in the figure legends, recombinant mouse C5a (500ng/mouse; R&D Systems, Minneapolis, MN, USA), and recombinant mouse CXCL4 (500ng/mouse; R&D Systems, Minneapolis, MN, USA). Sham treated mice received an equal volume of PBS i.t. alone. At the end of experiments (8-12 hours), mice were euthanized for collection of broncho-alveolar lavage fluids (BALF). BALF were centrifuged to separate cells before further analysis or cryopreservation at -80°C.

### Colorimetric assays

The CXCL4 ELISA kit was purchased from R&D Systems. The cell-free supernatants or cell-free broncho-alveolar lavage fluids were analyzed according to the manufacturer’s protocol. In short, samples were diluted in 0.1% BSA in PBS to fit into the standard range. Optical densities of oxidized TMB were measured with an Opsys MR Dynex microplate reader or a Tecan Infinite M Nano plate reader. Mouse albumin was quantified by ELISA (Bethyl Laboratories, Montgomery, TX, USA). Total protein was determined by BCA assay (Pierce, Rockford, IL, USA).

### Flow cytometry

For surface staining, the cells were washed in ice-cold sterile PBS (650g, 5 min, 4°C) and stained for 30 min with fixable viability dye eFluor 780 (Thermo Fisher Scientific, 1:1000) using heat-killed (1 min, 65°C) cells as positive controls. Next, cells were washed twice with FACS buffer [1% (w/v) BSA, 0.01% (w/v) sodium azide, 2.5 mM EDTA in sterile PBS], preincubated for 20 min with anti-mouse CD16/CD32 antibody (TruStain FcX, BioLegend, San Diego, CA, USA, 1:50) in FACS buffer followed by 20 min incubation with fluorescent dye-conjugated, anti-mouse antibodies for CD11b (clone M1/70, 1:100), Ly6G (clone 1A8, 1:50), F4/80 (clone BM8, 1:100), CD11c (clone N418, 1:600), SiglecF (clone E50-2440, 1:400), or matched fluorochrome-labeled isotype antibodies (all from BioLegend).

For phospho-flow cytometry, cells were washed with FACS buffer after stimulation, fixed for 20 min with Cytofix (BD Biosciences), washed with FACS buffer, re-suspended in pre-cooled (−20°C) Perm III buffer (BD Biosciences) and incubated overnight at -20°C. Thereafter, samples were washed again with FACS buffer, incubated for 15 min on ice with anti-CD16/CD32 blocking antibody (clone 93, BioLegend, 1:50) before antibodies against phospho-Akt(T308) (clone J1-223.371, BD Biosciences, 1:10), CD11b and F4/80 antibodies were added for additional 30 min. The cells were washed with Perm/Wash and re-suspended in FACS buffer.

At least 50,000 events of interest were acquired on a FACSCanto II or LSR II (BD Bioscience). For cell counting, 123count eBeads (Thermo Fisher Scientific, 1:15) were added according to the manufacturer’s instructions. FlowJo V10.0-V10.8.1 Software (FlowJo, Ashland, OR) was used for data analysis.

### Reverse transcription and real-time PCR

Macrophages were lysed and total RNA was isolated using the RNeasy Mini Kit (Qiagen, Venlo, Netherlands). Reverse transcription of cDNA was performed with the High-Capacity cDNA Reverse Transcription kit (Life Technologies, Carlsbad, CA) in a Mastercycler pro S (Eppendorf, Hamburg, Germany). Quantitative PCR was performed on a C1000 with CFX real time PCR detection system (Bio-Rad Laboratories) using 1-2 ng cDNA per sample complemented with iQ SYBR®Green Mastermix (BioRad) and specific forward and reverse primers at a concentration of 0.5 µM each. *Cxcl4* gene expression was compared between samples by normalizing to *Gaph* expression and applying the 2^-ΔΔCt^ formula (20). Primer sequences were as follows: mouse *Gadph* 5’-TACCCCCAATGTGTCCGTCGTG-3’ (forward), 5’-CCTTCAGTGGGCCCTCAGATGC-3’ (reverse); mouse *Cxcl4* (NM_019932) 5’-CAGCTAAGATCTCCATCGCTTT-3’ (forward), 5’-AGTCCTGAGCTGCTGCTTCT-3’ (reverse) (21).

### Histology

Lungs were inflated with 500 μl of paraformaldehyde (4%) for fixation, and 3 mm paraffin-embedded sections were stained with hematoxylin and eosin (H&E) in the histology core facility of the University Medical Center Mainz. An Olympus IX73 inverted oil immersion microscope with SC30 camera and Cell Sens Dimension V1 software was used for image acquisition (Olympus, Hamburg, Germany).

### Bead-based phosphoprotein assay

The phosphorylation status of Akt was evaluated in lysed macrophage samples (Bio-Plex Cell Lysis Kit; Bio-Rad) using the phospho-Akt(Thr308) kit from Bio-Rad following the manufacturer’s protocol. Phospho-Akt content reflecting fluorescence was measured in a Luminex-200/BioPlex-200 system (Bio-Rad) and normalized to fluorescence intensities of unstimulated control macrophages.

### Reagents

Polyphosphates were a gift from James H. Morrissey and Stephanie A. Smith. The polyphosphates included short-chain polyphosphates (S-PolyP) with a polymer length of 25 – 125 mer inorganic monophosphate residues [P_i_], medium-chain polyphosphates (M-PolyP) with a polymer length of 150 – 325 mer P_i_, and long-chain polyphosphates (L-PolyP) with a polymer length from 200 –1300+ mer P_i_. Lipopolysaccharide (Escherichia coli, serotype O111:B4) was purchased from Sigma-Aldrich. Wortmannin and LY294002 were obtained from Tocris Bioscience (Bristol, United Kingdom).

Recombinant PPX1-Fc and PPX1-D127N-Fc fusion proteins were expressed in *E. coli* as a customized project (Biologics International Corp, Indianapolis, IN, USA). Proteins were lyophilized after extensive dialysis against 1-fold PBS (pH7.4), and reconstituted in sterilized water before use. The purity was >90% as judged by SDS-PAGE analysis and endotoxin was <1 EU/μg as determined by gel clot endotoxin assay. Protein sequences are available as Supplementary File 1.

### Human single cell RNA-sequencing data analysis

The expression of CXCL4 and its receptor CXCR3 were assessed in an open access human lung transcriptome dataset to understand their cellular distribution. The human lung transcriptome dataset from Travaglini et al. (22) was comprised of normal uninvolved lung tissue from three patients undergoing lobectomy for lung cancer, which was sequenced after enzymatic digestion. The processed data from 10X Chromium sequencing with cell type (free) annotations was retrieved from cellxgene (23), re-normalized, and finally integrated using the Harmony (24) wrapper function in Seurat (25).

### Statistical analysis

Statistical analysis and graphs were prepared in Prism v7-9 (GraphPad Software). Data in graphs are depicted as mean ± standard error of the mean (S.E.M.). *In vitro* experiments were repeated as 2-3 independent biological replicates and are shown as representative or pooled data. *In vivo* experiments were done with the numbers of mice per group as indicated by symbols (circles, squares, triangles) in the figures. Two-group single comparisons were made with a two-tailed, unpaired Student’s t-test. Multiple comparisons were made with a one-way or two-way ANOVA. P values <0.05 were considered significant.

## Results

### Long-chain polyphosphates induce CXCL4 in macrophages via PI3K-Akt signaling

To study the role of polyphosphates in inducing CXCL4 release, cultures of bone-marrow derived macrophages (BMDMs) from C57BL/6J wild type mice were incubated for 24 h with long-chain (L-PolyP; mean polymer length of ∼700 inorganic monophosphate residues [P_i_]) or short-chain polyphosphates (S-PolyP; ∼70 P_i_). Long-chain polyphosphate-treated macrophage supernatants showed a substantial increase in CXCL4 concentrations compared to controls (resting macrophages), whereas short-chain polyphosphates had no observable effect (**Figure 1A**). Next, we investigated the kinetics of polyphosphate-mediated induction of CXCL4. Long-chain polyphosphates enhanced CXCL4 release in a dose- and time-dependent fashion from BMDMs and peritoneal macrophages (PEMs), respectively, while short-chain polyphosphates had no significant impact across the different concentrations (0.1 µM–1000 µM) and time points (0–48 h) tested (**Figure 1B, 1C**).

**FIGURE 1.**
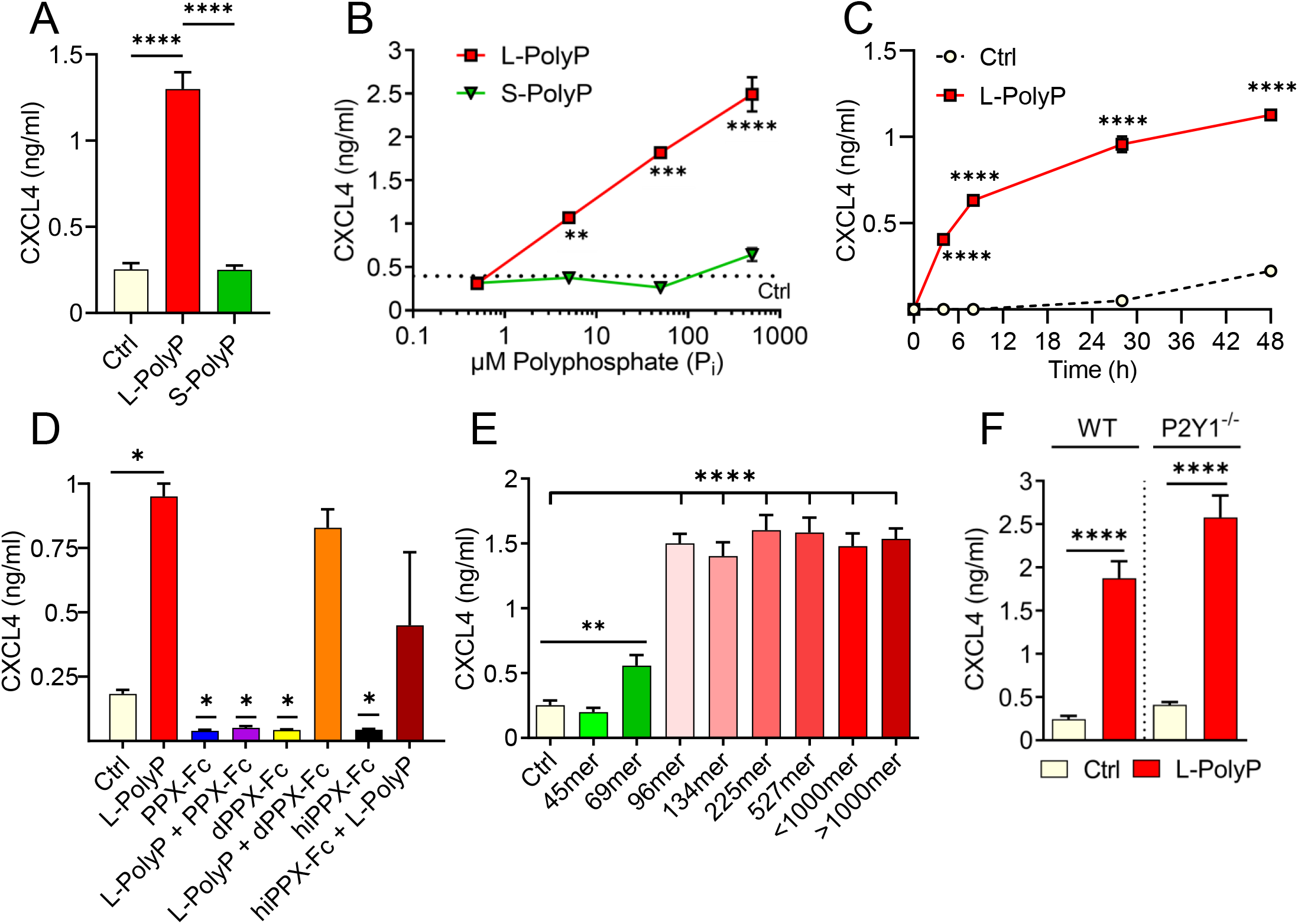
Polyphosphates induce CXCL4 release from macrophages. (**A**) CXCL4 in supernatants of bone marrow derived macrophages (BMDMs) from C57BL/6J wild type mice after incubation with long-chain polyphosphates (L-PolyP; P_i_700, 50 μM) or short-chain polyphosphates (S-PolyP; P_i_70, 50 μM) for 24 h. Resting macrophages served as controls (Ctrl). (**B**) Dose-response of CXCL4 release from macrophages (BMDMs) with different concentrations of long-chain or short-chain polyphosphates compared to basal levels from control macrophages, 24 h. (**C**) Time course of CXCL4 release from peritoneal elicited macrophages (PEMs) after long-chain polyphosphates (50 μM). (**D**) Long-chain polyphosphates (50 μM) were incubated overnight at 37°C with recombinant exopolyphosphatase-Fc fusion protein (PPX-Fc), mutated/dead exopolyphosphatase-Fc protein (dPPX-Fc), or heat inactivated exopolyphosphatase-Fc protein (hiPPX-Fc) followed by transfer to macrophages (PEMs) and CXCL4 detection after 24 h. (**E**) Polyphosphate polymers of narrow chain length distributions were incubated with macrophages (PEMs) followed by CXCL4 detection after 24 hours. (**F**) CXCL4 induced by long-chain polyphosphates in macrophages (BMDMs) from wild type mice compared to P2Y1^-/-^ mice. CXCL4 was measured by ELISA for all experiments shown. Polyphosphate concentrations were 50 μM in all experiments except for frame B. All data are representative of 3 independent experiments. Data are presented as mean ± SEM; *p < 0.05; **p < 0.01; ***p < 0.001; ****p < 0.0001.

The amounts of polyphosphates are expressed as concentrations of monophosphate units, meaning that 50 μM of long-chain polyphosphates (P_i_ 700) contain 10-fold fewer chains than 50 μM of short-chain polyphosphates (P_i_ 70). Based on our findings, 50 µM concentrations of polyphosphates at 24 h were considered as optimal conditions of strongest effect size and used for subsequent experiments with macrophages. No qualitative differences in the responsiveness to long-chain polyphosphates of either BMDMs or PEMs were noted (data not shown). Polyphosphates alone did not appear to generally induce other cytokines/chemokines that were tested (e.g., CCL2, CCL7, CCL9, CXCL10; data not shown) (8).

The long-chain polyphosphate effect on CXCL4 release was validated by incubating polyphosphates overnight at 37°C with recombinant exopolyphosphatase-Fc fusion protein (PPX-Fc), mutated/dead exopolyphosphatase-Fc protein (dPPX-Fc), or heat inactivated exopolyphosphatase-Fc protein (hiPPX-Fc), which were then transferred to macrophages for 24 h. As predicted, the activity of long-chain polyphosphates disappeared after exopolyphosphatase digestion. However, long-chain polyphosphates continued to induce CXCL4 in macrophages treated with non-functional exopolyphosphatases (dPPX-Fc, hiPPX-Fc; **Figure 1D**), confirming that polyphosphates have a direct and chain-length dependent effect on CXCL4 release. In another series of experiments using polyphosphate preparations with a more narrow range of chain lengths, we observed that polymers of ∼100mer length onwards exert a maximal potency to induce CXCL4 release in macrophages after 24 h (**Figure 1E**).

Interaction of polyphosphates with the P2Y1 receptor was reported to amplify inflammatory signals in endothelial cells (26). Therefore, we assessed whether P2Y1 also modulates the effects of polyphosphates on CXCL4 release in macrophages through an experimental design involving macrophages from wild type (WT) and P2Y1^-/-^ mice. P2Y1 did not appear to have any effect on long-chain polyphosphates-mediated CXCL4 release in macrophages (**Figure 1F**), suggesting that it is not required for the biological activity of polyphosphates in macrophages.

Toll-like receptor 4 (TLR4) is a key pattern recognition receptor that detects bacterial LPS and other PAMPs/DAMPs. TLR4 initiates inflammatory responses through the two adaptor molecules, MyD88 and TRIF, which are essential for all TLRs (27). Macrophages treated with a combination of long-chain polyphosphates and LPS showed a synergistic induction of CXCL4 compared with long-chain polyphosphates or LPS alone (**Figure 2A**). This synergy was also evident at the transcriptional level with higher CXCL4 mRNA abundance in macrophages treated with long-chain polyphosphates and LPS (**Figure 2B**). Next, we studied whether polyphosphates mediate CXCL4 induction also through TLR4. We examined the amount of CXCL4 released by polyphosphate- and LPS-treated macrophages from MyD88^-/-^ and TRIF^-/-^ mice side-by-side with wild type. Long-chain polyphosphates alone did not require MyD88 or TRIF for the induction of CXCL4, suggesting that polyphosphates are not ligands of TLRs (**Figure 2C**). There was a mild trend towards lower CXCL4 release in macrophages from MyD88^-/-^ mice treated with LPS alone or together with long-chain polyphosphates when compared to WT mice (**Figure 2C**). In contrast, TRIF^*-/-*^ macrophages seemed rather more responsive to the combination of LPS and long-chain polyphosphates. These findings fit with the concept that LPS activates CXCL4 gene expression mainly through the MyD88 pathway. The phosphatidylinositol 3-kinase-Akt serine/threonine kinase (PI3K-Akt) pathway is downstream of MyD88 and is involved in regulation of inflammatory responses (28). It is activated or suppressed based on phosphorylation of key amino acid residues of Akt (27). Macrophages treated with long-chain polyphosphates showed substantial phosphorylation of Akt at Threonine-308 (T308) in a time-dependent manner using a bead-based assay (**Figure 2D**). When macrophages were co-treated with either of two Akt inhibitors, Wortmannin or Ly294002, there was a marked reduction in the amount of CXCL4 released (**Figure 2E, 2F**). In another series of experiments, we observed that long-chain polyphosphates also augmented LPS-mediated phosphorylation of Akt at T308, as evident from the increased frequencies of CD11b^+^F4/80^+^ macrophages positive for p-Akt(T308) when compared to controls (controls: 1.57%; LPS: 14.1%; LPS + L-PolyP: 39.1%), as detected by flow cytometry (**Figure 2G, 2H**). These findings suggest that long-chain polyphosphates have a substantial impact on dysregulating immune responses by interacting with the PI3K/Akt pathway in macrophages.

**FIGURE 2.**
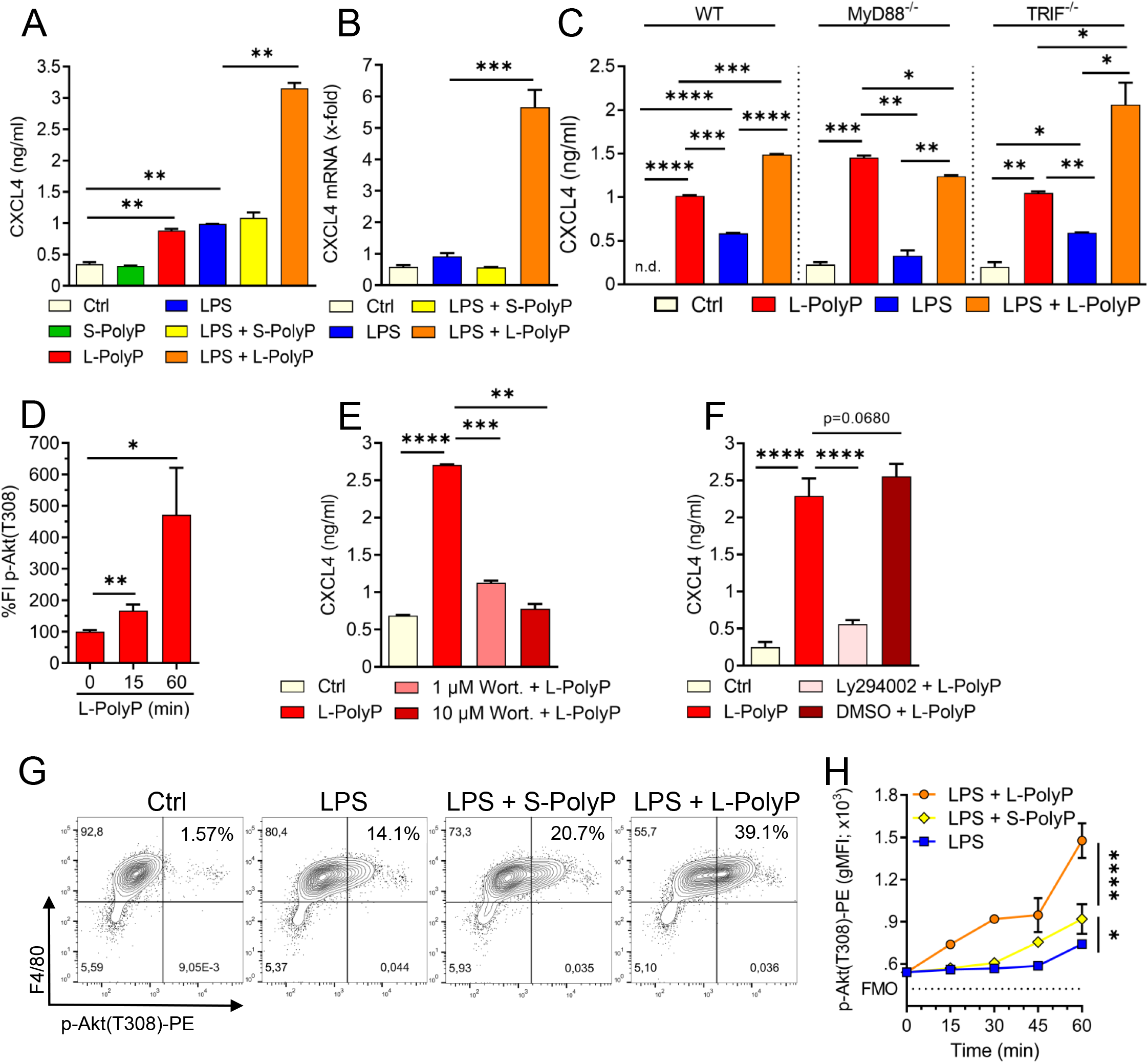
Polyphosphates regulate CXCL4 through PI3K/Akt signaling in macrophages. (**A**) CXCL4 release from C57BL/6J wild type macrophages (PEMs) after incubation with long-chain or short-chain polyphosphates ± LPS (100 ng/ml), Ctrl: resting control cells, 24h, ELISA. (**B**) CXCL4 mRNA expression in macrophages (PEMs) after polyphosphates ± LPS, 24h, RT-PCR. (**C**) CXCL4 release from macrophages (BMDMs) of wild type (WT), MyD88^-/-^ and TRIF^-/-^ mice, 24h, ELISA. (**D**) Relative quantification of phosphorylated Akt (threonine 308) in macrophages (BMDMs) at 0-60 min after long-chain polyphosphates. The values of fluorescence intensities (FI) were normalized to controls (0 min), bead-based assay. (**E**) CXCL4 release from polyphosphate-stimulated macrophages (BMDMs) co-treated with the Akt inhibitor, Wortmannin, 24h, ELISA. (**F**) CXCL4 release from macrophages (BMDMs) co-treated with the Akt inhibitor, Ly294002 (stock was dissolved in DMSO), 24h, ELISA. (**G**) Contour plots of phospho-Akt in F4/80^+^ macrophages (BMDMs) after activation with short/long-chain polyphosphates and LPS, 60 min, flow cytometry. (H) Geometric mean fluorescence intensities (gMFI) of pooled data (n=4/condition) as in frame G. Data are presented as mean ± SEM; *p < 0.05; **p < 0.01; ***p < 0.001; ****p < 0.0001: n.d.: not detectable.

### Long-chain polyphosphates promote lung injury

To investigate the specific contribution of polyphosphates to lung injury and inflammation, we injected polyphosphates into the trachea of C57BL/6J mice. Histopathological examination of mouse lungs after intra-tracheal long-chain polyphosphate administration showed diffuse hypercellularity caused by accumulation of PMNs, hyaline membranes, and alveolar wall thickening consistent with pulmonary edema and acute lung injury (**Figure 3A**). The total protein content in bronchoalveolar lavage fluid (BALF) was also significantly higher in long-chain polyphosphate-treated mice compared to sham controls (**Figure 3B**), consistent with greater disruption of the endothelial-epithelial barrier.

**FIGURE 3.**
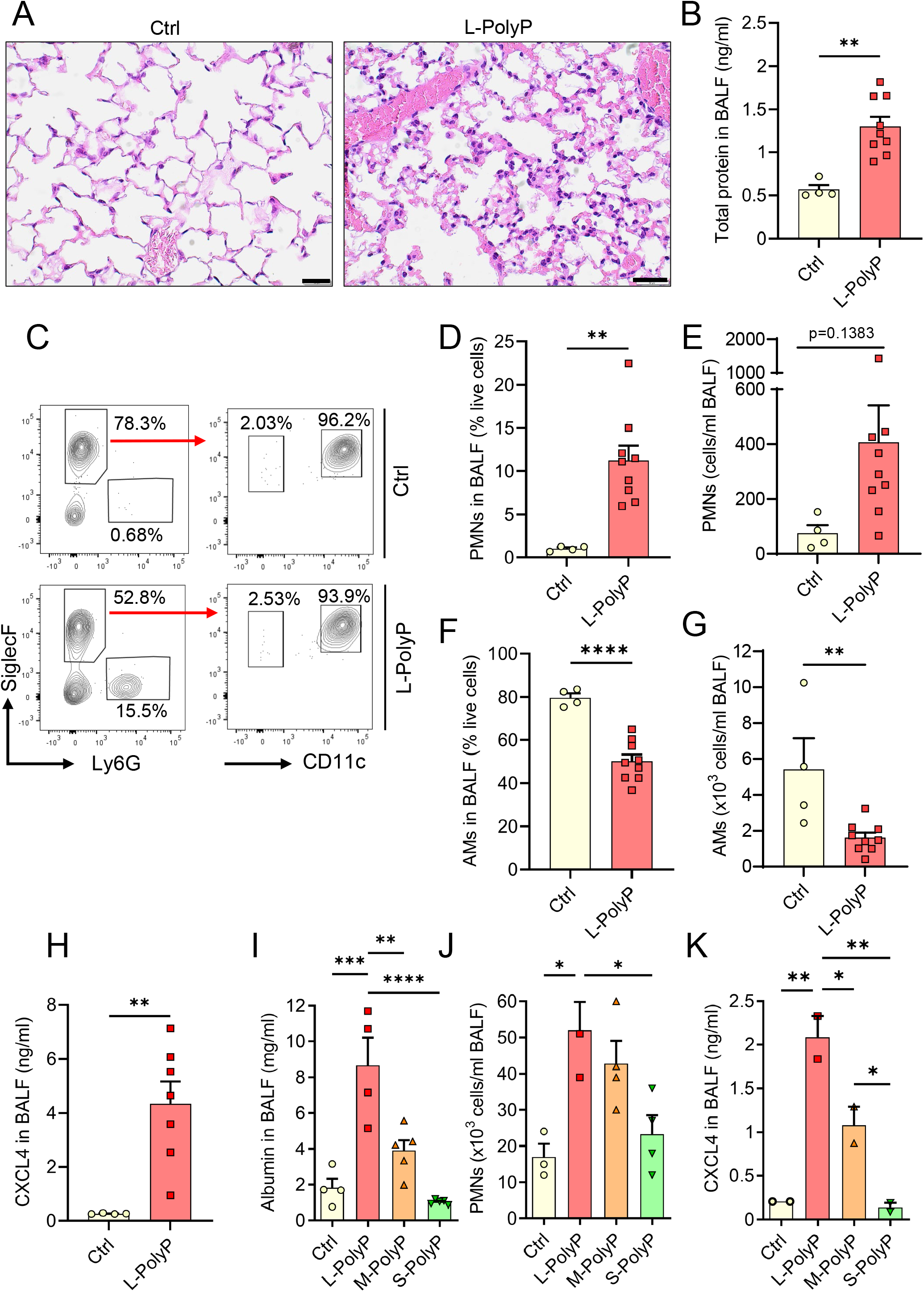
Long-chain polyphosphates cause lung injury. (**A**) Lung sections of C57BL/6J wild type mice obtained 8h after intra-tracheal (i.t.) administration of long-chain polyphosphates (40μl at 20 mM/mouse). Sham control mice (Ctrl) received PBS (40 μl/mouse, i.t.), H&E staining, scale bar: 20 μm. (**B**) Total protein in bronchoalveolar lavage fluids (BALF) after long-chain polyphosphates or sham control treatment, 8h, BCA assay. (**C**) Representative contour plots of Ly6G^+^ polymorphonuclear neutrophils (PMN), CD11c^+^SiglecF^+^ alveolar macrophages and SiglecF^+^Ly6G^-^CD11c^-^ eosinophils in BALF after polyphosphate-induced lung injury compared to sham controls, 8h, flow cytometry. (**D**-**E**) PMN frequencies and absolute numbers from mice as in frame C. (**F**-**G**) Frequencies and absolute numbers of alveolar macrophages (AMs) from mice as in frame C. (**H**) CXCL4 in BALF of mice in frame C. (**I**) Albumin in BALF of wild type mice administered i.t. with long-chain (L), medium-chain (M), short-chain (S) polyphosphates, or sham, 8h, ELISA. (**J**) PMN numbers in BALF from mice in frame I, manual count. (**K**) CXCL4 in BALF from C57BL/6J wild type mice as described in frame I. Data are presented as mean ± SEM; *p < 0.05; **p < 0.01; ***p < 0.001; ***p < 0.0001.

Immunophenotyping showed relatively higher influx of Ly6G^+^ PMNs and lower numbers of CD11c^+^SiglecF^+^ alveolar macrophages (AMs) in BALF after long-chain polyphosphate-induced lung injury compared to sham controls (**Figure 3C–3G**). Consistent with *in vitro* findings, there was augmented presence of CXCL4 compared to controls (**Figure 3H**).

Based on these observations, we next analyzed whether the extent of lung injury correlated with polyphosphate chain length. Long-chain polyphosphates consistently mediated significantly greater lung injury compared with medium-chain polyphosphates (M-PolyP; mean polymer length of ∼140 P_i_), short-chain polyphosphates, and controls, as demonstrated by the highest albumin content, highest influx of PMNs, and greatest induction of CXCL4 in BALF (**Figure 3J, 3K**).

We next sought to understand the role of CXCL4 in long-chain polyphosphate-mediated lung injury. As blocking antibodies and knockout mice for CXCL4 were not available, we performed studies testing recombinant CXCL4 administration. While recombinant CXCL4 alone caused a non-significant trend of PMN influx (4.2±0.6% vs. 1.0±0.1%) to the lungs compared to controls, we observed no synergistic effects with long-chain polyphosphates in terms of alveolar/endothelial barrier disruption (total protein in BALF) and numbers of alveolar macrophages and PMNs (**Supplementary Figure 1A**). When knockout mice for the promiscuous CXCL4 receptor, CXCR3, were studied, the only difference in the acute phase of polyphosphate-induced lung injury was a lower frequency of alveolar macrophages (**Supplementary Figure 1B**). Together, these data suggest a blank role for CXCL4 in polyphosphate-induced lung injury.

To extrapolate our findings to the human lung, we assessed the basal cellular expression of CXCL4 and CXCR3 in a publicly available normal lung single cell transcriptomic dataset. This analysis showed low basal mRNA expression of CXCL4 in macrophages. On the other hand, CXCR3 expression was only abundant in plasmacytoid dendritic cells (pDCs) and T cells (**Supplementary Figure 2**). Since T cells and pDCs are not many in number during the early phase of lung injury, this may explain some of the observed results.

### Cross-talk between long-chain polyphosphates and C5a in lung injury

Complement activation plays an important part in the host response to pathogens. Polyphosphates have been suggested to suppress the complement pathway (7, 29), although this mechanism has not been clearly elucidated. Elevated expression of the complement-derived anaphylatoxin, C5a, and interaction with its receptor, C5aR1, drive the pathophysiology of ALI (16, 17, 19). Here, we observed a significant increase in albumin content in BALF from mice that were co-administered both rmC5a and long-chain polyphosphates compared with each alone (**Figure 4A**). In addition, there was abundant release of CXCL4 in BALF from mice that received a combination of rmC5a and polyphosphates compared with mice that received only polyphosphates or rmC5a, and sham controls (**Figure 4B**). The effects of polyphosphates on C5a-mediated lung injury were partly explained by the elevated expression of C5aR1 on myeloid cells from long-chain polyphosphate-treated mice compared to controls (**Figure 4C**). The geometric mean fluorescence intensity (gMFI) values for C5aR1 were higher in alveolar macrophages and PMN from long-chain polyphosphate challenged mice compared to controls, though this difference was statistically significant only for alveolar macrophages (**Figure 4D, 4E**). The synergistic action of polyphosphates and C5a was further confirmed by studies using C5aR1^-/-^ mice, where we observed a significant reduction in albumin leakage and suppressed liberation of CXCL4 in long-chain polyphosphate-treated C5aR1^-/-^ mice compared with C57BL/6J wild type (WT) mice (**Figure 4F, 4G**). Collectively, these findings support that polyphosphate-mediated lung injury is amplified through a C5a-dependent mechanism.

**FIGURE 4.**
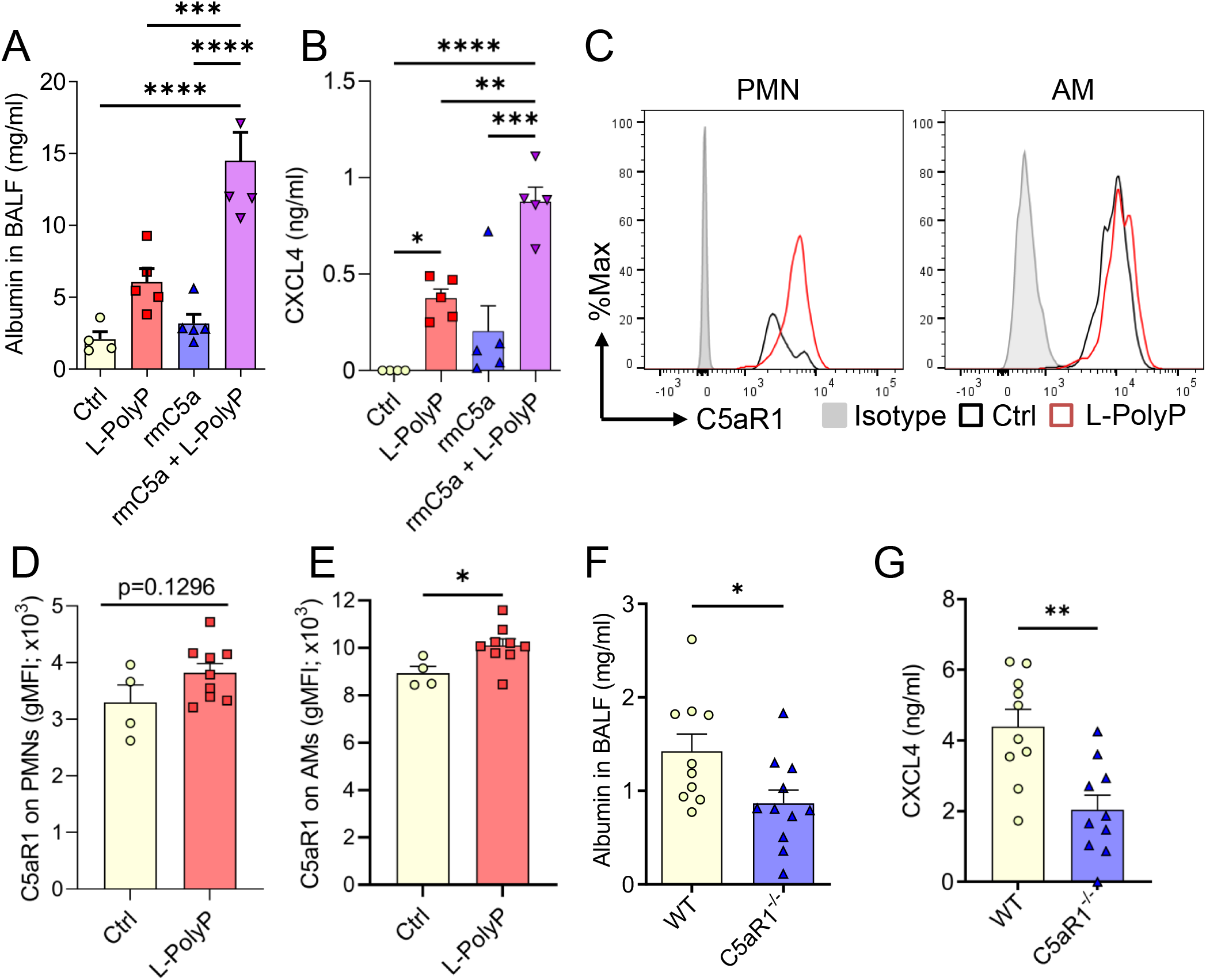
Polyphosphates synergize with C5a in lung injury. (**A**) Albumin in BALF of C57BL/6J wild type mice after long-chain polyphosphate-induced lung injury (40μl at 20 mM/mouse) ± recombinant mouse C5a (100 ng/mouse i.t.), 8h, ELISA. (**B**) CXCL4 in BALF from mice in frame A, ELISA. (**C**) C5aR1 histograms pre-gated on PMNs or alveolar macrophages (AMs) from mice after polyphosphate-induced lung injury or sham treated controls. (**D**-**E**) C5aR1 surface expression as geometric mean fluorescence intensities on PMNs and AMs from mice in frame C. (**F**) Alveolar albumin in wild type (WT) and C5aR1^-/-^ mice after long-chain polyphosphate-induced lung injury (40μl at 20 mM/mouse), 8h, ELISA. (**G**) CXCL4 in BALF from WT and C5aR1^-/-^ mice in frame F, 8h, ELISA. Data are presented as mean ± SEM; *p < 0.05; **p < 0.01; ***p < 0.001; ****p < 0.0001.

## Discussion

In this study, we have established that polyphosphates induce the typical signs of experimental lung injury (30), when administered to the airways of healthy mice. Polyphosphate concentrations are elevated in the blood and extracellular fluids during bacterial infection (8), although it is unknown if such polyphosphates are actively secreted or passively released. Bacterial pneumonia is a major cause that precipitates the development of ALI/ARDS in clinical settings (31). Hence, polyphosphates could represent a pathogenic mechanism linking bacterial infection to the development of ALI/ARDS. In addition, complement activation and C5a-mediated dysregulation of the inflammatory response are culprits in the pathophysiology of ARDS (16, 17, 19, 32). Elevated C5a is found in the lung epithelial lining fluid of ARDS patients, which resolves after recovery (33). In septic primates, anti-C5a antibodies protect against decreased oxygenation, pulmonary edema, and lethality (34). Our findings suggest that polyphosphates aggravate the influx of PMNs and destruction of the lung barrier caused by C5a (Fig. 4), whereas no synergistic effect of the polyphosphates and CXCL4 combination was observed for these endpoints (Supp. Fig. 1).

We have identified that the pathophysiological potency of polyphosphates increases with chain length, which demonstrates the distinctive roles of short-chain polyphosphates from platelets and long-chain polyphosphates from bacteria. In this study, we have denoted polyphosphate concentrations based on the amount of monophosphate residues, which is common practice in the field, given the limitations in quantitative methods to determine the precise chain length and the fact that polyphosphate preparations typically contain a range of chain lengths (8). This means that in our experiments, the number of polyphosphate chains is actually 10-fold higher in short-chain polyphosphates conditions as for long-chain polyphosphates. Notwithstanding, short-chain polyphosphates show very low to none biological activity for the studied endpoints. It is tempting to speculate that a chain length greater than P_i_60-70 could be needed because polyphosphates may simultaneously bind two proteins, which are then brought into spatial proximity by a long polyphosphate chain. It remains enigmatic whether bacterial polyphosphates are recognized by yet unidentified pattern-recognition sensors or if their biological activities are an integrated result of manifold interactions with several positively charged host proteins. Our finding that polyphosphates remained fully capable of inducing CXCL4 in MyD88^-/-^ and TRIF^-/-^ macrophages suggests that Toll-like receptors are not involved (Fig. 2C). We could not confirm a requirement for P2Y1 as a polyphosphate sensor in macrophages (8), although P2Y1 may transmit some effects in non-immune cells (26, 35).

While a classical ligand-receptor relationship remains unidentified, we uncover that polyphosphates activate PI3K/Akt signaling in macrophages. This expands earlier reports that polyphosphates recruit Akt and mTOR in human endothelial cells and breast cancer cells (5, 36). Moreover, we have shown that polyphosphates directly bind several human proteins associated with the PI3K/Akt pathway (37).

Polyphosphates mediate the release of their binding partner, CXCL4 (38), although both appeared not to synergize in fueling a more severe lung injury in our studies. Suggesting a dispensable role of CXCL4, we found that total protein leakage and PMN numbers in BALF are not different with co-administration of recombinant CXCL4 or in mice deficient in CXCR3. Of note, interpretation of findings in CXCR3^-/-^ mice is ambiguous because CXCL4, CXCL9, CXCL10 and CXCL11 all signal through CXCR3 (39). Even more complicating, CCR1 has recently been proposed as an alternative receptor for CXCL4 in a human monocytic cell line (40). Interestingly, CXCL4^-/-^ mice show improved histopathologic scores and lower total protein in BALF, albeit unchanged PMN numbers in acid aspiration-induced lung injury (41). Furthermore, CXCL4^-/-^ mice have diminished viral clearance and decreased PMN influx in the early stage of H1N1 influenza virus infection, which is followed by more severe lymphocytic lung pathology and higher mortality at later stages (42). In experimental bacterial pneumonia caused by *Pseudomonas aeruginosa*, CXCL4^-/-^ mice show defective bacterial clearance, more severe endothelial/epithelial permeability disturbances, reduced PMN accumulation in the lungs, and reduced blood platelet-neutrophil interactions (43). Transgenic mice overexpressing human CXCL4 are protected from early lethality (16-48 hours) after LPS-induced endotoxic shock (44). Thus, a pathophysiologic relevance of CXCL4 in infection and lung disease is supported by the literature, although we could not directly confirm this with our approach and at the early 8 hour time point studied.

In our experience, the activity profiles of polyphosphates are unique, specific and context dependent. Polyphosphates alone are not broadly proinflammatory or directly cytotoxic. In our laboratory, we routinely use polyphosphate-induced CXCL4 release as a positive control and convenient endpoint for validation in many studies. We propose the general concept that bacteria-derived, long-chain polyphosphates represent an immune evasion strategy which misdirects inflammation to cause tissue damage and prevent efficient pathogen clearance. Gram-negative bacteria encode genes for the enzyme, polyphosphatekinase (PPK), to synthesize polyphosphates from ATP. While gram-positive bacteria can also produce polyphosphates, no PPK homologs exist and alternate enzymatic machineries await identification. Instead of targeting polyphosphate synthesis in bacteria, we propose that administration of recombinant polyphosphatases could be of broad therapeutic value in sepsis and ALI/ARDS from both gram-negative and gram-positive bacteria. To further validate this potential approach, experimental studies of ARDS induced by infections with live bacteria will be needed in the future.

## Acknowledgments

This work was financed by the Federal Ministry of Education and Research (01EO1003, 01EO1503 to M.B.), the Deutsche Forschungsgemeinschaft (BO3482/3-3, BO3482/4-1 to M.B.) and the National Institutes of Health (R01AI153613, R01HL139641 to M.B.). C.R. was awarded a Fellowship of the Gutenberg Research College at the Johannes Gutenberg-University Mainz. We thank James H. Morrissey and Stephanie A. Smith for reagents and discussions. We thank Christian Gachet and Kristine Gampe for providing bone marrow from P2Y1^-/-^ mice. We thank Foruzandeh Samangan, Catherine O’Neal and Melissa Pesta for technical assistance. We cordially thank Archana Jayaraman for assistance with manuscript writing and data analysis. We thank Lucien Garo for reading the final version of the manuscript. The authors are responsible for the content of this publication.

## Author Contributions

J.R., S.W., A.S. and K.B. designed and performed experiments, and analyzed data. C.R. provided MyD88^-/-^ and TRIF^-/-^ mice. M.B. and J.R. wrote the manuscript, which was edited by S.W. and C.R. M.B. conceived and supervised the study, designed experiments, interpreted data, and provided funding.

