## Supplemental files for "Bacterial Polyphosphates Induce CXCL4 and Synergize with Complement Anaphylatoxin C5a in Lung Injury"

### Fc-PPX1 Protein Sequence:

Protein Length=635 MW=71833.0 Predicted pI=5.94

```
1   MEPKSCDKTH TCPPCPAPEL LGGPSVFLFP PKPKDTLMIS RTPEVTCVVV DVSHEDPEVK
61  FNWYVDGVEV HNAKTKPREE QYNSTYRVVS VLTVLHQDWL NGKEYKCKVS NKALPAPIEK
121 TISKAKGQPR EPQVYTLPPS RDELTKNQVS LTCLVKGFYP SDIAVEWESN GQPENNYKTT
181 PPVLDSDGSF FLYSKLTVDK SRWQQGNVFS CSVMHEALHN HYTQKSLSLS PGKDDDDKMS
241 PLRKTVPEFL AHLKSLPISK IASNDVLTIC VGNESADMDS IASAITYSYC QYIYNEGTYS
301 EEKKKGSFIV PIIDIPREDL SLRRDVMYVL EKLKIKEEEL FFIEDLKSLK QNVSQGTELN
361 SYLVDNNDTP KNLKNYIDNV VGIIDHHFDL QKHLDAEPRI VKVSGSCSSL VFNYWYEKLQ
421 GDREVVMNIA PLLMGAILID TSNMRRKVEE SDKLAIERCQ AVLSGAVNEV SAQGLEDSSE
481 FYKEIKSRKN DIKGFSVSDI LKKDYKQFNF QGKGHKGLEI GLSSIVKRMS WLFNEHGGEA
541 DFVNQCRRFQ AERGLDVLVL LTSWRKAGDS HRELVILGDS NVVRELIERV SDKLQLQLFG
601 GNLDGGVAMF KQLNVEATRK QVVPYLEEAY SNLEE
```

### Fc-PPX1-D127N Protein Sequence:

Protein Length=635 MW=71844.7 Predicted pI=5.81

```
1   MEPKSCDKTH TCPPCPAPEL LGGPSVFLFP PKPKDTLMIS RTPEVTCVVV DVSHEDPEVK
61  FNWYVDGVEV HNAKTKPREE QYNSTYRVVS VLTVLHQDWL NGKEYKCKVS NKALPAPIEK
121 TISKAKGQPR EPQVYTLPPS RDELTKNQVS LTCLVKGFYP SDIAVEWESN GQPENNYKTT
181 PPVLDSDGSF FLYSKLTVDK SRWQQGNVFS CSVMHEALHN HYTQKSLSLS PGKDDDDKMS
241 PLRKTVPEFL AHLKSLPISK IASNDVLTIC VGNESADMDS IASAITYSYC QYIYNEGTYS
301 EEKKKGSFIV PIIDIPREDL SLRRDVMYVL EKLKIKEEEL FFIEDLKSLK QNVSQGTELN
361 SYLVNNNDTP KNLKNYIDNV VGIIDHHFDL QKHLDAEPRI VKVSGSCSSL VFNYWYEKLQ
421 GDREVVMNIA PLLMGAILID TSNMRRKVEE SDKLAIERCQ AVLSGAVNEV SAQGLEDSSE
481 FYKEIKSRKN DIKGFSVSDI LKKDYKQFNF QGKGHKGLEI GLSSIVKRMS WLFNEHGGEA
541 DFVNQCRRFQ AERGLDVLVL LTSWRKAGDS HRELVILGDS NVVRELIERV SDKLQLQLFG
601 GNLDGGVAMF KQLNVEATRK QVVPYLEEAY SNLEE
```

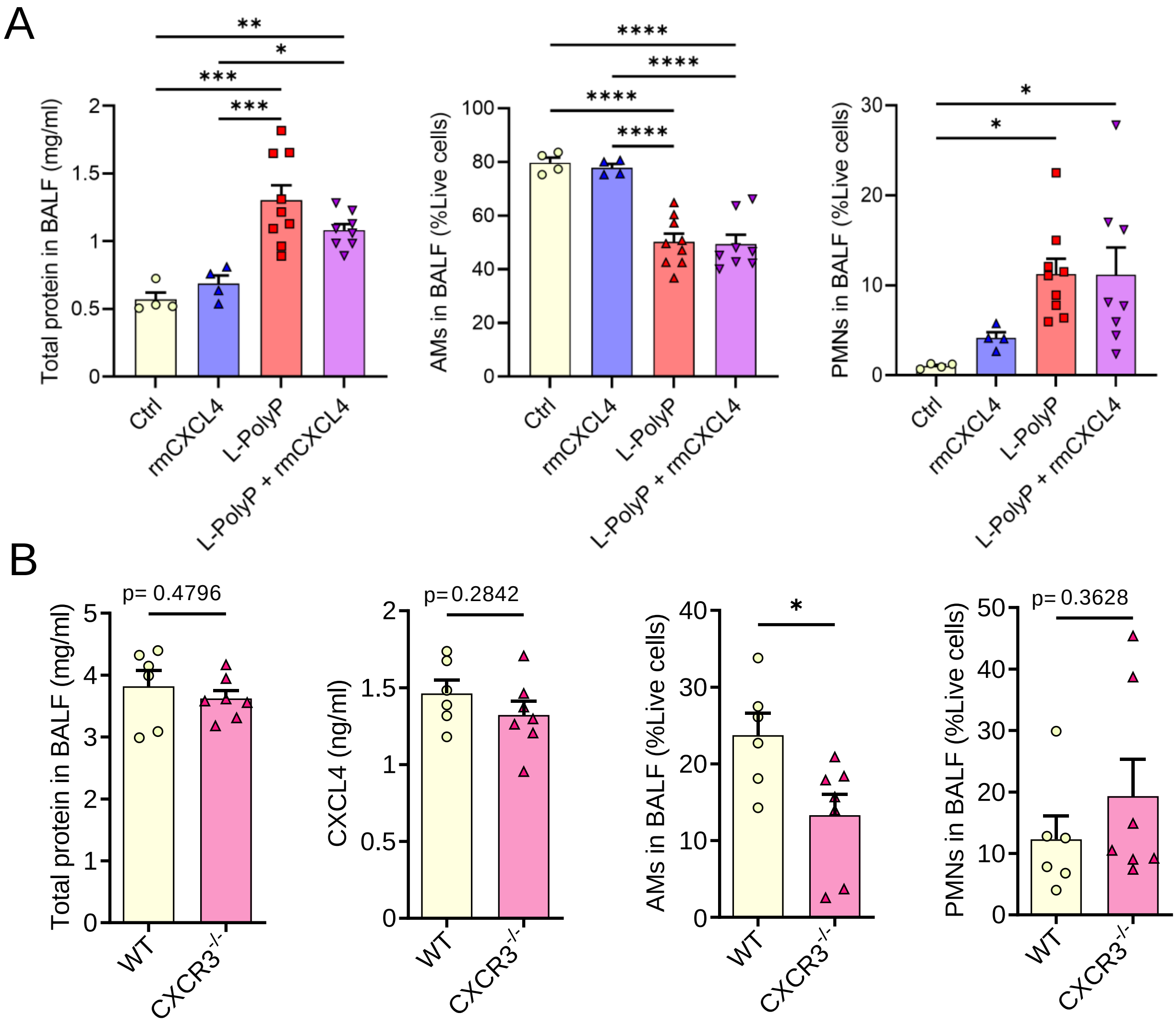


**SUPPLEMENTARY FIGURE 1. Role of CXCL4 and CXCR3 in polyphosphate-induced lung injury.** (**A**) C57BL/6J mice (WT) received recombinant mouse CXCL4 (rmCXCL4, 500 ng/mouse i.t., n=4), long-chain polyphosphates (40µl of 20 mM/mouse i.t., n=9) alone or in combination (n=8). The sham control mice (Ctrl) received buffer (40µl PBS i.t., n=4). Total protein and frequencies of live CD11c$^{+}$SiglecF$^{+}$ alveolar macrophages (AMs) and live Ly6G$^{+}$ polymorphonuclear neutrophils (PMNs) were quantified in BALF by BCA protein assay and flow cytometry, respectively, 8h. (**B**) C57BL/6J and CXCR3$^{-/-}$ mice received long-chain polyphosphates (40µl of 10 mM/mouse i.t.). Total proteins, CXCL4 (ELISA), frequencies of live CD11c$^{+}$SiglecF$^{+}$ AMs, live Ly6G$^{+}$ PMNs were quantified in BALF (n=6-7/group), 12h. Data are presented as mean ± SEM; *p < 0.05; **p < 0.01; ***p < 0.001; ***p < 0.0001.

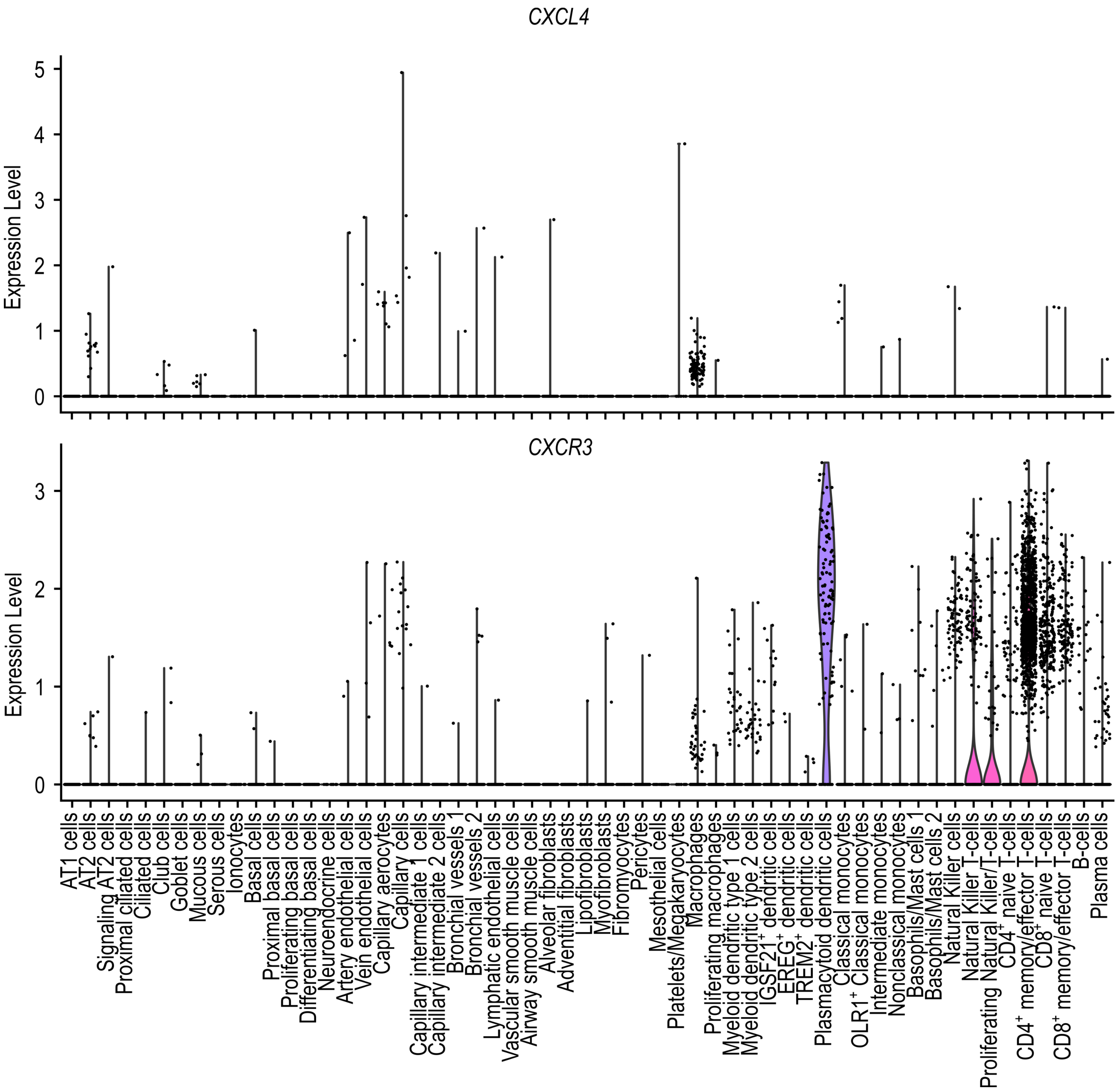


**SUPPLEMENTARY FIGURE 2. Expression of *CXCL4* (*PF4*) and *CXCR3* in normal human lung transcriptomes.** Violin plot showing expression levels from single cell RNA-sequencing data from adult human lungs (n=3). The data is based on scRNA-sequencing by Travaglini et al. from healthy, uninvolved lung tissues from patients (aged 46 years [male], 51 years [female], 75 years [male]) undergoing lobectomy for pulmonary tumors.
